# Cycle length flexibility: is the duration of sexual receptivity associated with changes in the social environment?

**DOI:** 10.1101/2022.09.16.508123

**Authors:** Darmis Fragkiskos, Huchard Élise, Cowlishaw Guy, Carter J. Alecia

## Abstract

Research in social mammals has revealed the complexity of strategies females use in response to female-female reproductive competition and sexual conflict. One point at which competition and conflict manifests acutely is during sexual receptivity, indicated by swellings in some primates. Whether females can adjust their sexual receptivity from cycle to cycle to decrease reproductive competition and sexual conflict in response to changes in the social environment has not been tested. As a first step, this study explores whether sexual receptivity duration is predicted by the social environment in wild female chacma baboons (*Papio ursinus*). Given that female baboons face intense reproductive competition and sexual coercion, we predicted that: females could shorten the duration of their sexual receptive period to reduce female-female aggression and male coercion or increase it to access multiple or their preferred male(s). We quantified 157 ovulatory cycles from 46 wild females living in central Namibia recorded over 15 years. We found no support for our hypothesis; however, we found a negative correlation between maximal-swelling duration and group size, a proxy of within-group competition. This study provides further evidence that swelling is costly for females as well as a testable framework for future investigations of ‘cycle length manipulation’.

## Introduction

Recent advances in sexual selection theory underline the importance of female-female competition, along with mate choice, in generating variance in female reproductive success [1–4]. In the first case, females tend to face competition with consexuals over access to males and reproductive resources [5–8]. This competition for access to males increases when there is synchrony of females’ fertility (i.e., ovarian synchrony), which creates a female-biased operational sex ratio (OSR) and increases aggression received by receptive females [9,10]. In general, females compete with consexuals for (a) their preferred mates (e.g., chacma baboons *Papio Ursinus*: [9]; red-fronted lemurs *Eulemur rufifrons*: [11]), (b) paternal services for their offspring (e.g., chacma baboons: [12,13]) and (c) other reproductive resources [14], such as breeding territories (e.g., lek center [15]) or food resources [16]. Those benefits are achieved either through monopolisation of resources directly from females (i.e., contest competition for food) [17] or indirectly (e.g., nutrient-rich spermatophores in insects [18], low predation risk in lek centers [15]). Antagonism with consexuals can manifest through non-aggressive interactions (e.g., signals [18]) and escalate up to physical aggression, which can be detrimental to the receiver’s short-or long-term fitness [19].

Considering mate choice, it may be advantageous for females to concentrate paternity into a single male who can provide benefits in the form of good genes or paternal services such as protection to her or her offspring. However, in group living species, a number of social pressures may decrease the possibility for females to express their mating preference. For example, recent studies highlight the role of sexual conflict [20], which frequently manifests via infanticide in mammals. Indeed, infanticide risk may push females to mate with multiple rather than single partners, to dilute paternity certainty among multiple males who subsequently restrain from attacking an offspring that they could have sired [21,22]. Female choice may be further constrained by sexual coercion, when males direct aggression to females [3,20,23], either to increase the probability that they will mate with them or to decrease the probability that they will mate promiscuously [24]. In such cases, females will mate with their aggressors if resisting is more costly than giving in [20]. In groups of water striders (*Aquarius remigis*), aggressive males acquire more matings than less aggressive ones [25]. In primates, male baboons (*Papio ursinus*) that attack pre-ovulatory females are more likely to monopolize them when receptive [26] and females do not copulate close to adult males to avoid bystanders’ aggression [26,27]. Also, female chimpanzees (*Pan troglodytes*) initiate copulations most frequently with those males that were most aggressive towards them throughout their cycle [28,29] and male lemurs (*Lemur catta*) that are not preferred can force copulations [30]. However, male coercion can lead to female counterstrategies and antagonistic co-evolution between the sexes [20] since coercion is associated with multiple costs for females. For example, the social dynamics involving male immigration events, high male-male competition and unstable male hierarchies are stressful for females because they typically increase the intensity of coercion [13,31,32] and harassment [33–35]. Moreover, females suffer costs from coercion when: they are forced to mate with subordinate or incompatible males; direct aggression or harassment increases stress hormone secretion [29]; males reduce their preferred level of parental care [23]; or coercion impairs female choice [31].

By decreasing the length of their receptive period, females could potentially decrease (1) time that they are subject to aggression they receive from other females (female-female competition) (2) and/or males (sexual coercion). By contrast, by increasing the length of their receptive period, females could potentially (3) increase their probability of mating with a preferred male (female choice) or (4) multiple males (paternity confusion). Evidence for manipulation of female cycles exists [19,35–37], with studies detecting physiological changes in female cycling patterns (e.g., changes to cycle length, resumption of cycling) as stress responses to the social environment. Socially-stressed primate females, for example by being harassed near menstruation, have longer menstrual cycles [35]. Cycling female yellow baboons (*Papio cynocephalus*) that receive aggression from consexuals tend to have more cycles to conception and longer interbirth intervals [19], while pregnant females that are systematically harassed experience reduced fitness (through abnormal gestation length, spontaneous abortion, and premature delivery) [19]. Psychosocial stress stemming from social instability, in the form of male take-overs, also induces female physiological responses: the presence of unfamiliar males can mediate early sexual maturation [38,39] or abortion [40,41] in female mammals, in a likely effort to limit infanticide risk. Novel males in geladas (*Theropithecus gelada*) prompt females of any reproductive state (immature, cycling, lactating, or pregnant) to start or resume cycling [39,42], while female hamadryas (*Papio hamadryas*) with young infants develop swellings to advertise a reduced postpartum amenorrhea that results in copulations with novel males during take-overs [37]. However, these swellings are deceptive signals that ultimately decrease infanticide risk (four out of five infants survived in [37]) since females did not conceive during the first cycles after the take-over, their reproduction was not accelerated and the interbirth intervals were not shorter compared to times with no male take-overs. Despite the above results, no study to date has proposed a theoretical framework that considers cycle length manipulation as a counterstrategy to sexual selection pressures. Additionally, the aforementioned studies provide evidence of cycle manipulation as an acute reaction to social pressures but have not explicitly tested or explored the fluctuations in sexual receptivity in response to changes in females’ social environment.

In this study, we explored whether there is a correlation between females’ oestrous duration and the social environment. We assumed that females could adjust their oestrous cycles to either reduce aggression from (1) coercive males or (2) females or to increase their access to (3) their preferred or (4) multiple males. We focused on female chacma baboons because they exhibit elevated female-female competition compared to other primates [13,43,44], live in multimale-multifemale groups, mate promiscuously [26,45], and have exaggerated swellings that reliably indicate females’ stage of the follicular phase of the oestrus cycle [36]. Female chacma baboons are an ideal model to study female-female competition, intersexual competition, as well as sexual coercion, since they target the dominant male as a sexual partner [13,44] but also mate promiscuously [9] outside their peak swelling to establish more males with a nonzero paternity probability or to initiate male-male competition (e.g., sperm competition) [27,36]. In addition, males’ monopolisation of only one receptive female at a time (i.e., mate-guarding: [7,12]) results in female-female competition over sexual access to particular males [9,13], who might be better able to protect them from aggressive consexuals or provide future postpartum care for the offspring [46].

To address our overarching aim, we proceeded in two steps. First, we explored the variation within and among females in the durations of swelling (SD) and maximal-swelling (MSD) stages of the cycle and we predicted that different mechanisms could act on each part of the cycle. We predicted that there would be more variation within females’ cycles than between females across each of their cycles. Second, we tested whether variation in the duration of specific cycle phases was associated with four aspects of the social environment (summarised in Table 1).

**Table 1:**
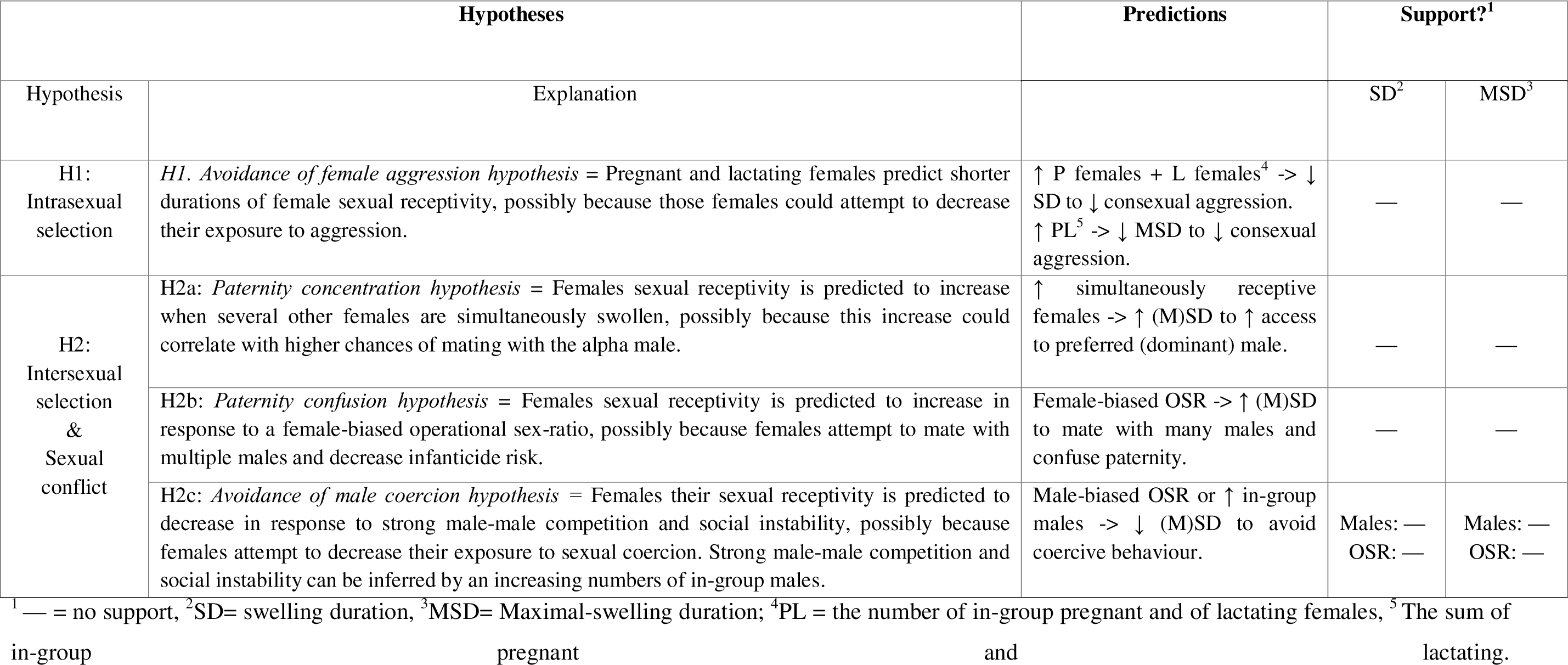
Hypotheses, predictions and support for the hypotheses proposed to affect the duration of female cycle-specific phases.

First, considering female intrasexual selection, we tested whether female competition for paternal care was associated with variation in the duration of receptivity (the avoidance of female aggression hypothesis [H1]). Pregnant and lactating females aggressively target swollen females to decrease competition for paternal care of their own offspring, or to induce reproductive suppression so that their own infants are not born at the same time as many others [33]. For this reason, we predicted a positive correlation between in-group pregnant and lactating females and aggression in cycling females that would result in receptive females terminating the swelling earlier and, ultimately, decreasing their exposure to consexual aggression.

Our second hypothesis dealt with intersexual selection and sexual conflict (H2): we hypothesized that females’ SD and MSD would show a correlation with competition for access the dominant mate (the paternity concentration hypothesis, H2a). We predicted that, when competition for access to the alpha male was likely because there were simultaneously-swollen individuals, females would respond by delaying ovulation—through an elongation of their swelling or MSD—to increase the probability of mating with this male. By doing so females may benefit from (a) good genes [13,36] and (b) concentrating paternity certainty in their preferred male, usually the dominant, who is also the most likely to pose a future risk of infanticide if they fail to mate with him [13,47].

We also tested whether the risk of infanticide from multiple males correlated with patterns of variation in female baboons’ cycle-phases (the paternity confusion hypothesis, H2b). We predicted that an increase in swelling duration (SD) and maximal-swelling duration (MSD) will be associated with a female-skewed operational sex ratio (OSR) so that females can mate with many males to reduce infanticide risk. Finally, we tested whether the effect of social instability in the form of high numbers of in-group males (i.e., strong male-male competition) correlated with patterns of variation in a female’s receptivity phases. We predicted that a male-skewed OSR or a high number of in-group males translates into high male-male competition, which in turn increases the risk of sexual coercion and male-initiated aggression for receptive females [12,26]. Under this scenario, a negative association between oestrous duration and the number of males or a male-biased OSR might provide evidence for females’ attempts to escape or minimize the aggression received from coercive males (the avoidance of male coercion hypothesis, H2c).

## Materials and Methods

### Study site and animals

Data were collected from identifiable females within two habituated groups, J and L, of a wild chacma baboon population at Tsaobis Leopard Park (22°22’S, 15° 44’E) in central Namibia. Studies on this population, which have been ongoing since 2000 [48], have revealed that the naturally-foraging baboons [48] exhibit low predation risk [49] and high female-female aggression rates [9,13] that can result in reproductive suppression [13]. Over the course of the study, troop numbers fluctuated around a median of 55 (44–69) in J troop and 52 (21–71) in L troop.

The data used in this study were collected periodically over 15 years, from 2005 to 2019, during field seasons lasting between 2 and 8 months *per annum*, usually during the austral winter. During field seasons, troops were visited daily, when possible, with some interruptions for other tasks or unforeseen circumstances. During troop visits, we collected data on troop composition (see below) and individual behaviour *ad libitum*.

Individuals’ ages were known from observing births or from patterns of dentition wear following capture. Individuals’ relative ranks were calculated annually from *ad libitum* recorded dominance interactions in the package Matman 1.1.4 (Noldus Information Technology, 2013). Absolute ranks were converted to a relative scale ranging from 0 (lowest rank) to 1 (highest rank), to control for group size, using the formula *1-((1-r)/(1-n))*, with *r* being individual’s absolute rank ranging from 1 to the total group size *n*. Group size was calculated as the total number of individuals in the troop for a particular year. Individuals who were present for less than half of a field season (due to emigration or death) were not included.

### Troop composition and reproductive state data

Each day that the troop was contacted, a census was completed that recorded the identities of the individuals present and the reproductive states of each adult female. Females’ reproductive states were recorded as: (1) pregnant (determined *post hoc* based on lack of resumption of swelling, reddening of the paracallosal skin and subsequent birth); (2) lactating (i.e., period following the birth of an infant until cycle resumption); (3) oestrous/swollen (i.e., exhibiting periovulatory swelling of the anogenital region); (4) non-swollen (i.e., deturgescent but not pregnant). For females in oestrus, we also recorded swelling sizes on a semi-quantitative scale from 0 (smallest) to 4 (largest), in order to capture within-individual variation in swelling size across successful oestrus cycles [50]. As mentioned above, baboon troops were not followed daily, and as such there were some missing data during some females’ cycles. We discarded any observations of the entire oestrous cycle where the start or end of the swelling duration was missing and thus could introduce inaccuracy.

For females in oestrus, swelling duration was calculated as the number of continuous days that a female was recorded with a swelling of any size. Because different mechanisms could act on the different parts of the cycle, in addition to swelling durations, we calculated the MSD as the number of days the focal female exhibited the largest swelling size during that respective cycle. In summary, we used 157 receptivity observations for swelling duration from 46 females (median number of swellings / female = 5, range = 1-14) and 150 receptivity observations for maximal-swelling duration (median number of swellings / female = 5, range = 1-14). The difference between the number of observations of these two variables stems from the fact that for seven observations of swelling duration the start and end date of receptivity were accurately known, yet this was not the case for the maximal-swelling duration of those cycles.

We calculated eight predictor variables describing aspects of the social environment: to consider (H1) female-female competition for paternal care, we calculated (1) the number of pregnant and (2) of lactating females, as well as (3) their sum (PL), during the time a focal female was started swelling. To consider (H2a) competition for females’ access to the preferred male, for the first day of a focal female’s swelling we calculated the number of simultaneously swollen females that were (4) not maximally-swollen and (5) maximally-swollen. To consider sexual conflict in terms of (H2b) infanticide risk and (H2c) sexual coercion, we calculated the (6) operational sex ratio (OSR) as the ratio of sexually active females (i.e., swollen at any level) to adult males and the (7) the number of in-group adult males. Finally, to account for food and resource competition we included group size in our models (8) if adding this covariate improved model performance.

### Statistical analyses

All statistical analyses were performed using R (version 4.0.3, 2020-10-10).

### Question 1: Repeatability of receptivity

To determine how much variation in females’ receptivity part of the cycle was due to differences between individuals, we calculated the repeatability (R) of the SD and MSD of all the females that had at least two measurements for each response (n = 38). R is calculated as 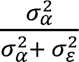, where 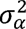 represents the between-group variance, 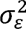 the within-group variance (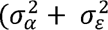 = the total phenotypic variance, *V_p_* [51]). A relatively high R estimate would indicate that the dependent variable exhibits high between- and low within-female variance; low values of R indicate traits with high within-female or low between-female variation [51]. We calculated R using the `rpt` function (`rptR` package), which can calculate repeatability for Poisson-distributed data, and the models included female identity and troop as random effects. Finally, both models controlled for the possible confounding fixed effects [52] of age and rank. The permutations and bootstrapping were set to 1000 and confidence intervals at 95%. However, we are aware that the repeatability we present in the results might be somewhat flawed as our response variables are overdispersed (the `rptR` package offers no way to correct for that excess variance though).

### Question 2: Social determinants of oestrous length

To determine whether the social environment was correlated with females’ fluctuations in sexual receptivity duration, we used two generalised linear mixed models (GLMMs; `glmer` function, `lme4` package [53]) with the response variables: swelling duration (SD) and maximal-swelling duration (MSD). Both SD and MSD were modelled using a negative binomial distribution to correct for overdispersion. Female identity nested in troop identity was included as a random effect for SD models and female identity was a random effect for MSD models, since variation between troops was absent for MSD. In both models, we included rank and age as control variables. To control for possible type I errors [54], we tested whether a model with random slopes for each of our predictors performed better than one without (Supp. Mat. Tables A2 & A3), using the corrected AIC criterion, AICc to compare models (`aictab` function, `AICcmodavg` package [55]). We included this analysis as some authors argue that random slopes are crucial to avoid false positive rates [56,57]. We also performed full-null model comparisons using a likelihood-ratio test. The null models contained only the control variables age and relative rank and, as random factors, female ID nested in troop for SD and female ID for MSD (see above). In both cases, the full models performed better than the null model (Supp. Mat. Table A4), so we proceeded with these full models for our analysis. Residuals and Q-Q plots were also checked for normality (for model diagnostics see Supp. Mat. Figures A1-A4).

We used variables to test our hypotheses. Firstly, we determined, with the use of AICc (as above), whether group size—the number of adults and sub-adults present in a troop—should be included as a control variable in the models and which predictors should be used to test for the effect of pregnant and lactating consexuals on sexual receptivity duration (H1). For group size, larger groups could result increased competition for limited resources (i.e., feeding competition), which could impact females’ condition and their swelling duration [7]; while for H1 we did not have an *a priori* prediction of whether the number of pregnant and of lactating females or the combined number (i.e., sum) of pregnant and lactating females (PL) is a better predictor. The results of model selection led us to include pregnant and lactating females separately for SD (but not group size), and the sum, as well as group size, for MSD (Supp. Mat. Table A1). In both SD and MSD models, we included an interaction with rank because rank usually determines who can harass whom. To explore the effect of the number of receptive females on sexual receptivity duration (H2a), we used the number of not maximally-swollen females when a focal female started swelling when the response was SD; and the number of maximally-swollen females when a focal female started swelling when the response was MSD. Thirdly, to determine if the risk of infanticide (H2b) or male-male competition (part of H2c) predicts fluctuations in female sexual receptivity durations, we included the OSR as a predictor. Lastly, to determine the response of sexual receptivity durations to male-male competition (part of H2c), we included the number of in-group males because it is a proxy of social instability in the sense that male ranks are more volatile when there is more competition (i.e., more males) (A. Carter, E. Huchard, pers. obs.). Lastly, we tested whether group size should be included as a control variable.

The best-fit model for SD and MSD, carrying 97% and 69% of the cumulative model weight (AICcWt), respectively, and showing the lowest AICc values, did not include random slopes. For the two resulting models we checked for multicollinearity of model terms using variance inflation factors (VIFs; Supp. Mat. Figures A2 & A4 respectively; `check_collinearity` function, `performance` package [58], a VIF < 5 indicates a low correlation of the predictor with other predictors and a VIF > 10 a high correlation [59]). All VIFs were < 5, except interaction terms, which is expected [60], so these variables were retained.

## Results

Females were swollen for an average (median) of 22 days (range 8-44) and maximally-swollen for a median of 6 days (range 1-24). There were no differences between troops (Mann-Whitney U test: SD, W = 3481.5, p = 0.10; MSD, W = 2651.5, p = 0.59).

### Question 1: Repeatability of receptivity

Neither SD nor MSD were repeatable at the individual level (SD: R = 0.036, SE = 0.049, CI = [0, 0.167], p = 0.251 & MSD: R = 0.054, SE = 0.058, CI = [0, 0.2], p = 0.283) (Supp. Mat. Table A5; Figure A5).

### Question 2: Social determinants of SD & MSD

We included two control variables in both our models, while for MSD we also included group size as a predictor of feeding and contest competition (Table 2). Rank was not significantly associated with either of our responses (SD: β = 0.25, SE = 0.26, p > 0.05; MSD: β = 1.02, SE = 0.62, p > 0.05; Figure 1), and neither was age (for SD, β = −0.01, SE = 0.004, p = 0.069; for MSD, β = −0.02, SE = 0.01, p = 0.056; Figure 1). Group size, on the other hand, predicted a decrease in MSD (β = −0.18, SE = 0.07, p = 0.013; Figure 1; Figure 2; Table 2). Specifically, a one-unit increase in group size negatively correlated with a drop of almost 16% in MSD (approximately 1.22 fewer days of MSD for each additional group member).

**Figure 1:**
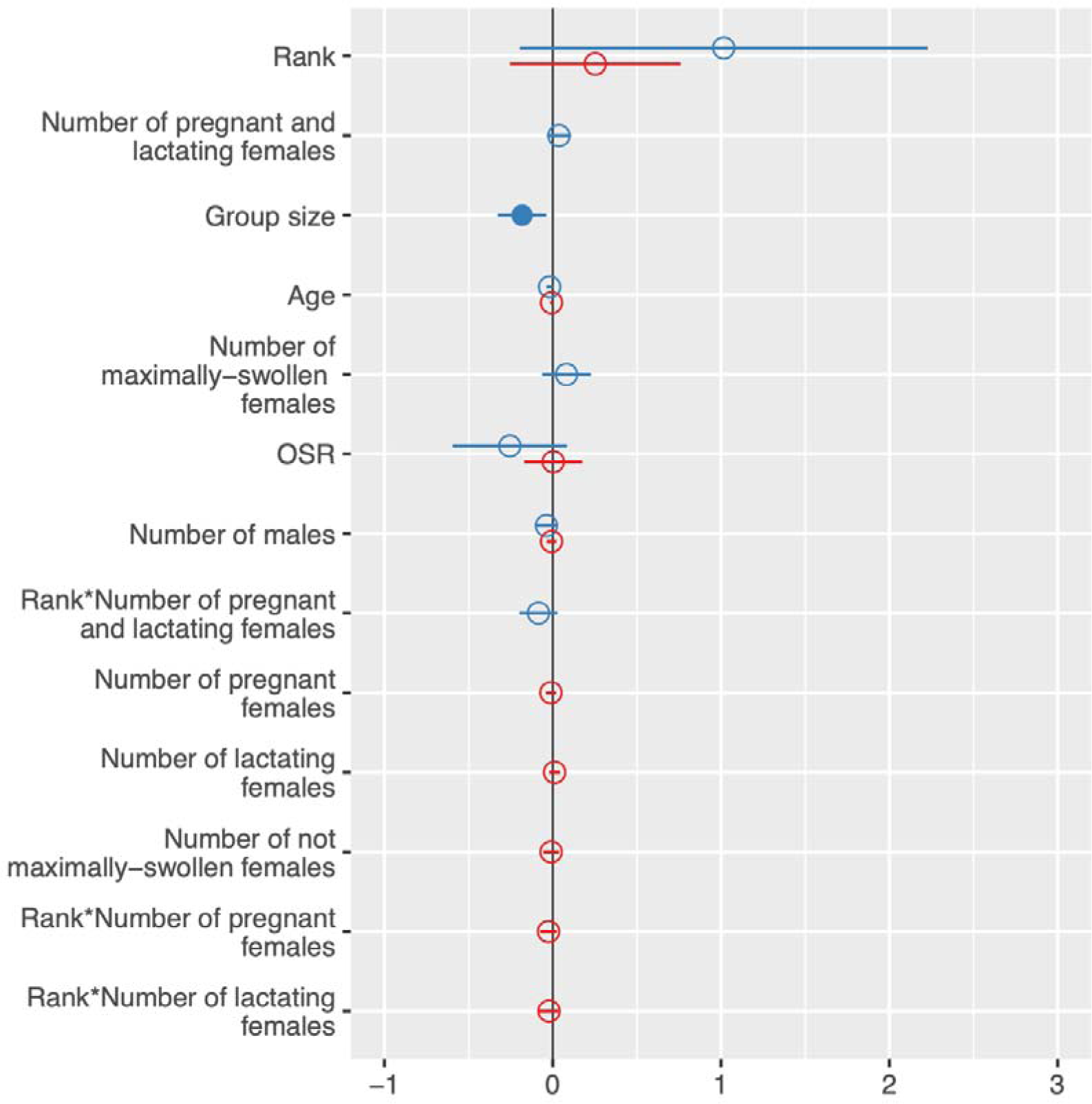
Forest plot of standardised estimates of the models with swelling duration (red) and maximal-swelling duration (blue) as response variables. The level of significance is indicated by an unfilled circle (p > 0.05) or a filled circle (p < 0.05). Only group size significantly predicted a decreased maximal-swelling duration.

**Figure 2:**
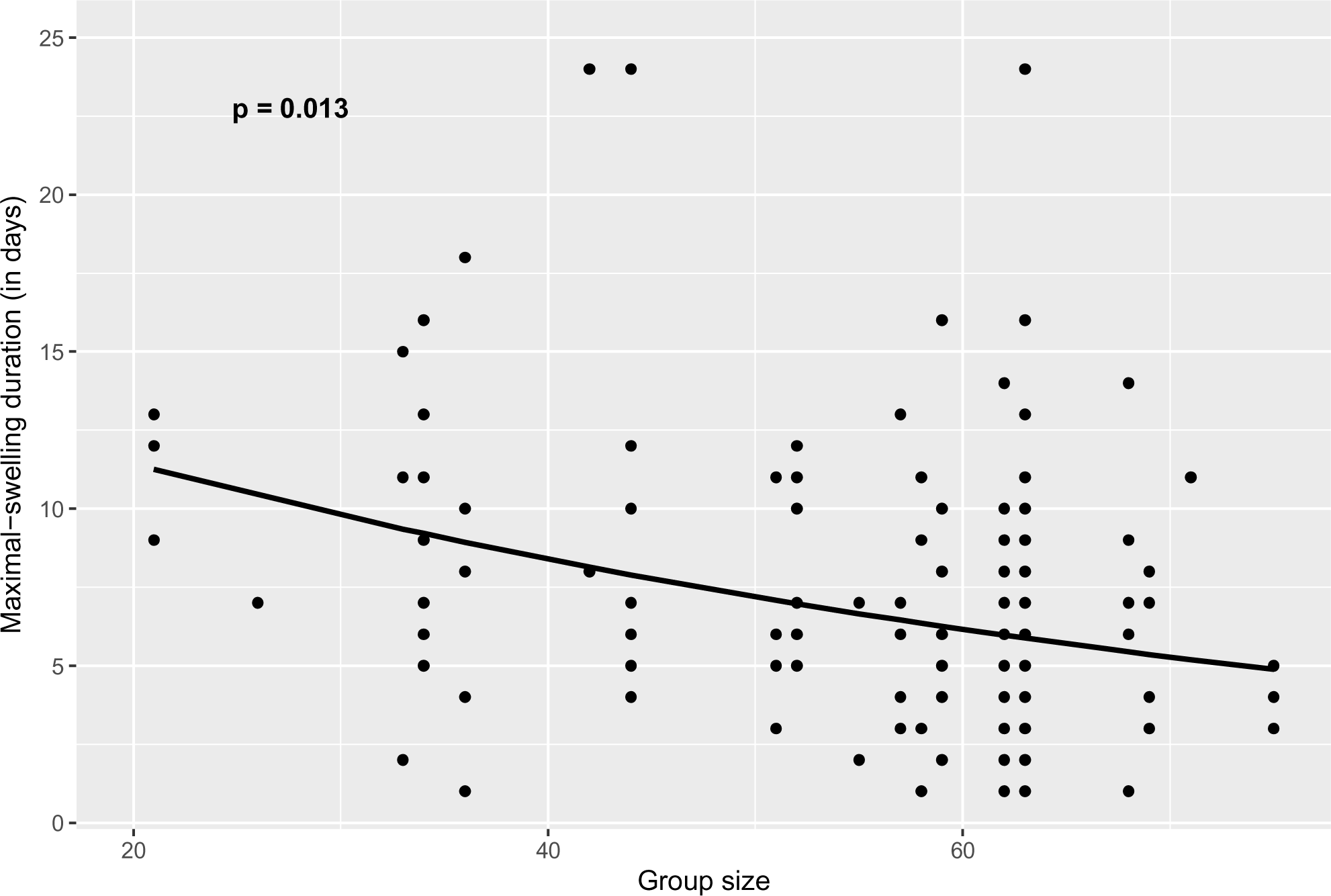
The effect of group size on maximal-swelling duration. The line indicates the predicted effect by varying the respective focal variable and by setting all the other covariates to their mean. The shaded area represents the 95% CI.

**Table 2:** Results of the two models with SD and MSD as response variables. “Log-Mean” measures the effect of the specific predictor on the response variable and is given in log-scale. “SE” indicates the standard error and CI shows the 95% confidence intervals. Significant results are indicated with bold.

| *Coefficients* | **Model 1 (SD)** |  |  |  | **Model 2 (MSD)** |  |  |  |
| --- | --- | --- | --- | --- | --- | --- | --- | --- |
|  | *Log-mean*<br>*of*<br>*estimates* | *SE* | *CI (95%)* | *p-value* | *Log-mean*<br>*of*<br>*estimates* | *SE* | *CI (95%)* | *p-value* |
| Intercept | 3.28 | 0.19 | 2.90 – 3.66 | **<0.001** | 2.00 | 0.55 | 0.92 – 3.07 | **<0.001** |
| Rank | 0.25 | 0.26 | -0.26 – 0.76 | 0.330 | 1.02 | 0.62 | -0.20 – 2.23 | 0.100 |
| Number of pregnant and lactating females | — | — | — | — | 0.04 | 0.04 | -0.04 – 0.11 | 0.314 |
| Rank*Number of pregnant and lactating females | — | — | — | — | -0.08 | 0.06 | -0.20 – 0.03 | 0.143 |
| Rank*Number of pregnant females | -0.03 | 0.03 | -0.07 – 0.02 | 0.317 | — | — | — | — |
| Rank*Number of lactating females | -0.02 | 0.03 | -0.08 – 0.03 | 0.444 | — | — | — | — |
| Number of lactating females | 0.01 | 0.02 | -0.02 – 0.04 | 0.524 | — | — | — | — |
| Number of pregnant females | -0.01 | 0.02 | -0.04 – 0.02 | 0.483 | — | — | — | — |
| Age | -0.01 | 0.004 | -0.02 – 0.00 | 0.069 | -0.02 | 0.01 | -0.04 – 0.00 | 0.056 |
| Number of not maximally-swollen ♀ | -0.01 | 0.02 | -0.06 – 0.04 | 0.681 | — | — | — | — |
| Number of maximally-swollen ♀ | — | — | — | — | 0.08 | 0.07 | -0.06 – 0.23 | 0.271 |
| Number of ♂ | -0.01 | 0.02 | -0.04 – 0.02 | 0.627 | -0.04 | 0.03 | -0.10 – 0.03 | 0.261 |
| OSR (♀/♂) | 0.003 | 0.09 | -0.17 – 0.18 | 0.976 | -0.26 | 0.17 | -0.60 – 0.08 | 0.141 |
| Group size | — | — | — | — | -0.18 | 0.07 | -0.33 – -0.04 | **0.013** |

### H1: Intrasexual selection

#### Avoidance of female aggression hypothesis

Under our first hypothesis, we tested whether the number of pregnant and lactating females was associated negatively with females’ SD or MSD. There were on average 10 (median, range = 3-28) pregnant and lactating females (PL), 5 pregnant (median, range = 0-11) as well as 6 lactating (median, range = 0-18) during females’ swelling and maximal-swelling phase. There was no relationship between the latter two variables and SD (pregnant: β = −0.01, SE = 0.02, p > 0.05; lactating: β = 0.01, SE = 0.02, p > 0.05) or between PL and MSD (β = 0.04, SE = 0.04, p > 0.05; summary of statistics at Table 2; Figure 1). There was also no effect of the interaction of these variables with rank.

### H2: Intersexual selection and sexual conflict

#### H2a Paternity concentration hypothesis

To test our second hypothesis, we tested whether there was positive association between SD and MSD and the number of swollen females. There were on average 2 (median, range = 0-6) not maximally-swollen females and 0 (median, range = 0-3) maximally-swollen females when a focal female started swelling. Neither response was associated with swelling durations (SD, not-maximally-swollen females: β = −0.01, SE = 0.02, p > 0.05; MSD, maximally-swollen females: β = 0.08, SE = 0.07, p > 0.05; Table 2; Figure 1).

#### H2b Paternity confusion hypothesis

We predicted a positive correlation between females’ SD or MSD and a female-biased OSR. The mean OSR value was 0.64 (range 0.1–2.5): on average there were more sexually active males than females. Our analyses showed no correlation between OSR and SD (β = 0.003, SE = 0.09, p > 0.05) or MSD (β = −0.26, SE = 0.17, p > 0.05; Table 2; Figure 1).

#### H2c Avoidance of male coercion hypothesis

We tested whether there was a negative correlation between the number of males and receptivity durations. There were on average 5 males (median, range = 1-10) when females started to swell. We found support for this hypothesis either (SD: β = −0.01, SE = 0.02, p > 0.05; MSD: β = −0.04, SE = 0.03, p > 0.05; Figure 1; Table 2).

## Discussion

The present study investigated the variation in the length of receptivity within versus across females. We provided a theoretical framework for why females could evolve to manipulate their receptivity—a mechanism we termed cycle length manipulation. We further tested whether variation in female sexual receptivity could be associated with changes in the social environment. We detected high within-female variation in the length of sexual receptivity. However, we found no evidence that variation in females’ sexual receptivity is strategically adjusted to decrease intra-or inter-sexual competition. We did, however, find that group size, and presumably within-group competition, negatively affected females’ maximal swelling durations.

We found that SD and MSD were not repeatable. Our results do not mean that there are no between-individual differences in SD and MSD, but that those differences are not large relative to within-individual differences and not consistent between females. Our findings show that cycle phases are labile traits that females could potentially manipulate. At a comparative level, high variation in oestrous duration has been detected in animals (e.g., captive jaguars, with a mean duration of 10.42 and a range of 7 to 15 days [61]). Women, in contrast, show repeatable cycle lengths, even though there is some variation within [62] and across women [63], as well as extensive variation in average cycle length across human populations [64]. It may be that baboons’ low repeatability of receptivity reflects the harsh environmental conditions at Tsaobis, with females’ condition fluctuating in response.

We found no evidence that the numbers of pregnant and lactating consexuals correlated with receptivity duration. One reason why our predictor did not affect receptivity duration could be that female reproductive suppression in the form of aggression was given by the female friends of a particular male [12]. Specifically, aggression was initiated by pregnant and lactating females who were friends of the male that was mate-guarding the receptive female. A more targeted approach that considers as a predictor the number of female friends a swollen female’s mate-guarding male has might provide evidence for cycle length manipulation. Alternatively, it may be that the particular stage of lactation and pregnancy might be a better predictor than just the total number of individuals in each phase. For example, a study in yellow baboons revealed an increasing attack rate initiated from lactating females as lactation stages progressed [19].

We predicted that higher numbers of receptive females would prompt an elongation of oestrous duration, so that females could concentrate paternity in the alpha male but found no support for this. At least two explanations may account for this absence of evidence. First, because female chacma baboons are non-seasonal breeders [13], oestrous synchrony in this population is low, with a median of two simultaneously swollen females (i.e., swollen at any phase of their cycle). Consequently, mating with the preferred male is possible for most cycling females and therefore elongation of the fertile period is not necessary. Second, the substantial aggression among cycling females in Tsaobis may limit females’ propensity to extend their receptivity period [9,13]. This is predominantly true for lower-ranking oestrous females and females who are not mate-guarded, which tend to be the usual target of aggression from higher-ranking females.

We found no support for hypothesis 2b, that there would be a positive association between an OSR with receptivity durations. Our results could suggest that paternity dilution through extended receptivity comes at a cost for females, which we believe is an increased rate of female-female competition and aggression. Under this scenario, it seems that females entering the mating pool are primarily aggressive towards consexuals at peak swelling, which are most likely monopolised by the dominant male.

The number of adult males, as well as the OSR, were not associated with receptivity durations, as suggested by H2c. While male aggression, intimidation and coercion are costly for females in this population [26,27], it could be that a decrease in the length of swelling might incur higher costs, such as a smaller swelling, which could deter higher-ranking males from mate-guarding. Alternatively, selection to elongate receptivity due to increased infanticide risk (seen at a comparative level across Cercopithecine primates [36]) but also to decrease receptivity to avoid coercion could result in no effect overall.

We found that group size correlated negatively with MSD. There are at least two possible explanations for this relationship that are not mutually exclusive and could operate simultaneously. Firstly, as group size increases, competition for limited resources intensifies and maintaining energetically-costly maximal swellings could become more difficult for females [7]. In addition, larger baboon groups tend to travel longer distances over greater areas, which could generate higher energetic demands [65]. This interpretation recognizes the energetic limitation that arises due to elevated levels of feeding competition. Secondly, in baboons, socially-induced stress may generate higher cortisol levels [66] and subsequently result in a decrease of the swollen period [33,35]. As the number of in-group individuals increases, receptive females might be subject to higher aggression rates, both from consexuals (intrasexual aggression) and heterosexuals (intersexual aggression). Female primates of larger groups tend to show higher fecal glucocorticoid concentrations (fGC) [65,67], which could decrease females’ receptivity durations indirectly. Under this scenario, social stress is interlinked with social competition, and future studies can directly test for that by including hormonal data and behavioural observations. Group size, however, was not a good predictor in the case of SD. This might be because (1) SD is comparatively less costly to maintain, even under stress, compared to MSD; or/and (2) competition with females and males is comparatively less during the whole SD compared to MSD.

In summary, this study correlated variation in the length of female sexual receptivity with aspects of the social environment that may influence the intensity of female reproductive competition and intersexual conflict. We found no support for sexual conflict or female-female competition driving receptivity duration. We did, however, find that one aspect of the social environment—group size, and presumably within-group competition—correlated with MSD in females. This suggests that maintaining large swellings is costly for females. Further investigations may benefit from including behavioral and hormonal data in order to integrate more closely the proximate mechanisms with ultimate explanations when testing this or related hypotheses.

## Ethics

Our research procedures were evaluated and approved by the Ethics Committee of the Zoological Society of London and the Ministry of Environment and Tourism, Namibia (MET Research/ Collecting Permits 886/2005, 1039/2006, 1186/2007, 1302/2008, 1379/2009, 1486/2010, 1486/2011, 1696/2012, 1786/2013, 1892/ 2014, 2009/2015, 2147/2016, 2303/2017, RPIV00392018/2019) and adhered to the ASAB/ABS Guidelines for the Treatment of Animals in Behavioural Research and Teaching.

## Data accessibility Authors’ contributions

AJC, EH and FD contributed to the conceptualization. AJC, EH designed the study. GC established the study system and oversaw data collection, with assistance from AJC and EH. FD performed the statistical analyses. All authors contributed to draft the manuscript. All authors gave final approval for publication and agreed to be held accountable for the work performed therein.

## Acknowledgments

We would like to thank the past Tsaobis Baboon Project leaders, students, volunteers and interns for contributing to the long-term data collection at the field site. This research was carried out with the permission of the Ministry of Environment and Tourism, the Ministry of Land Reform, and the National Commission on Research, Science, and Technology. We further thank the Tsaobis beneficiaries for permission to work at Tsaobis, the Gobabeb Namib Research Institute for affiliation, and Johan Venter and the Snyman and Wittreich families for permission to work on their land. Some findings formed part of a dissertation submitted in partial fulfilment of the requirements of the degree of MSc of the University College London in September 2021. This paper is a publication of the ZSL Institute of Zoology’s Tsaobis Baboon Project. Contribution ISEM n°202x-xx.

