## Supplementary material for "Cycle length flexibility: is the duration of sexual receptivity associated with changes in the social environment?"

**Table A1**

The models that were compared using AICc to determine, for each response variable, if group size should be used and if the sum of the number of pregnant and lactating females should be used as instead of the separate variables.

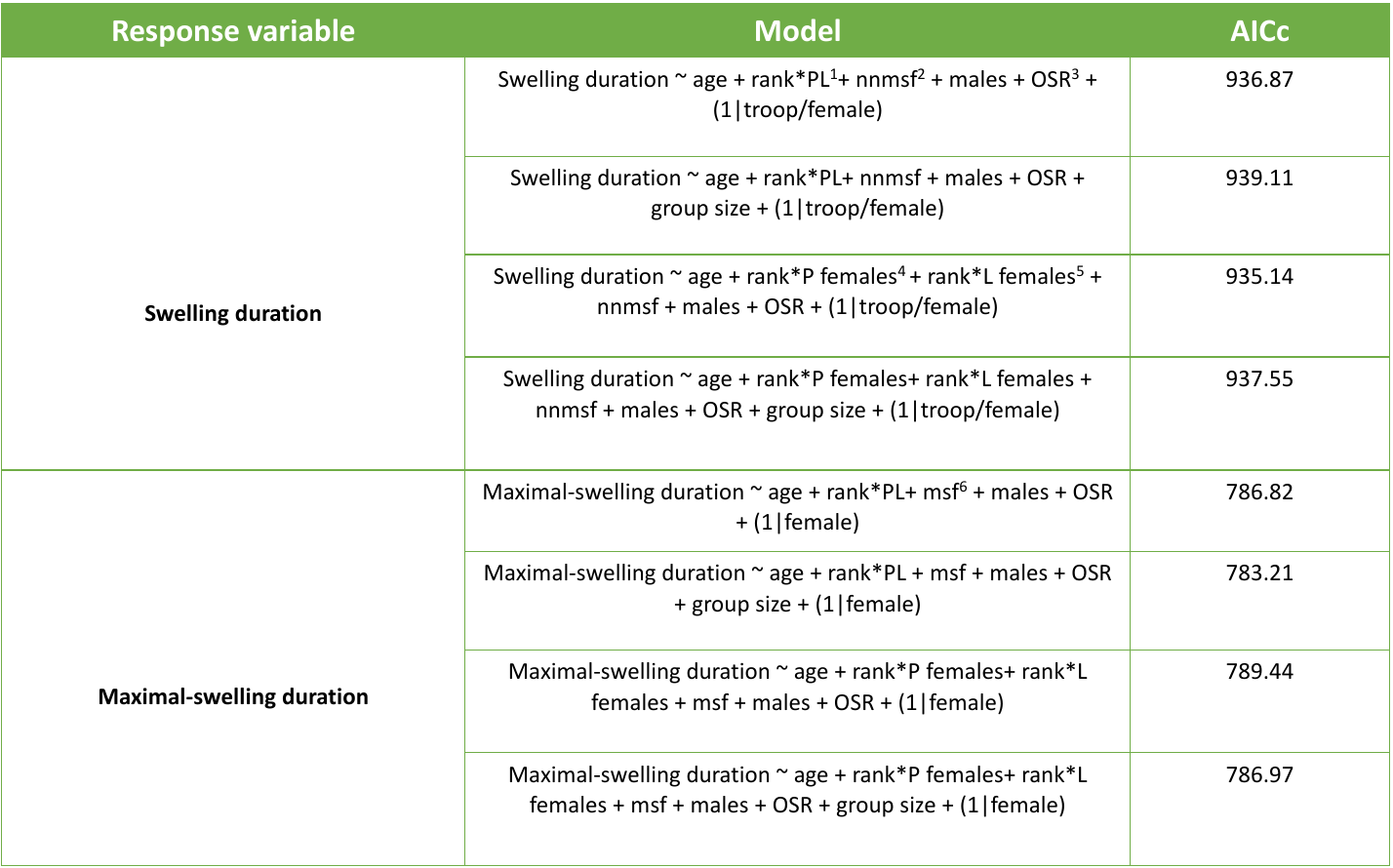

^1^Number of pregnant and lactating females specified with an interaction with rank; ^2^Number of not-maximally swollen females when the focal female started swelling; ^3^Operational sex ratio as the number of the number of sexually active females to adult males; ^4^Number of pregnant females interacting with rank; ^5^Number of lactating females interacting with rank; ^6^Number of maximally-swollen females when the focal females started swelling;

**Table A2**

The models (with swelling duration as the response variable) that were compared using AICc to determine if random slopes contribute to model fit. “Model problems” report any issue encountered when running the respective model (some of which might be false positives), and “AICc weight” indicates the cumulative model weight, that is it the proportion of the total amount of predictive power provided by all the models contained in each model. Here, the first model shows the lowest AICc value and carries the highest proportion of the cumulative model weight.

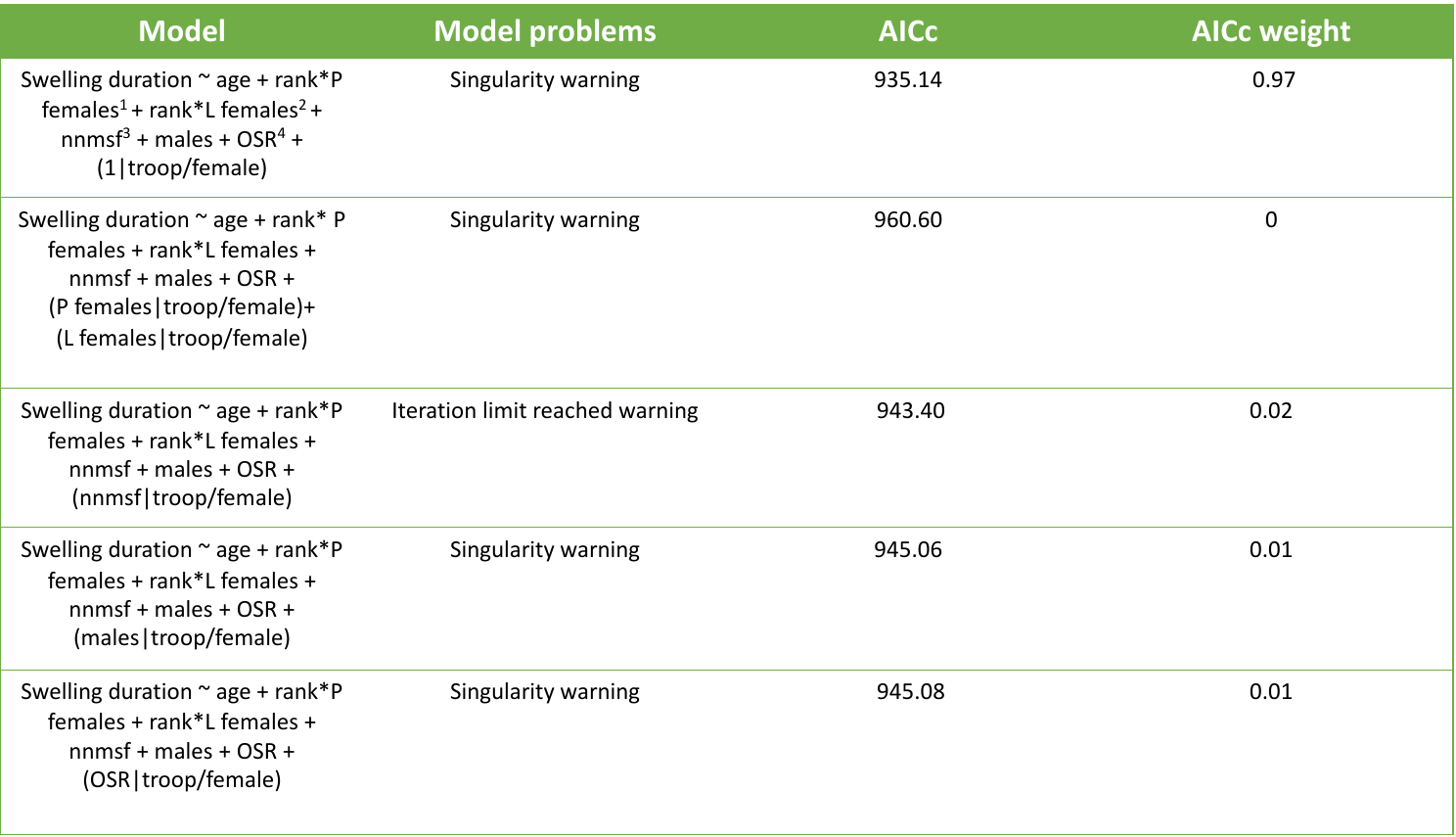

^1^Number of pregnant females with an interaction with rank; ^2^Number of lactating females with an interaction with rank; ^3^Number of not-maximally-swollen females when the focal started swelling; ^4^Operational sex-ratio as the number of the number of sexually active females to adult males.

**Table A3**

The models (with maximal-swelling duration as the response variable) that were compared using AICc to determine if random slopes contribute to model fit. “Model problems” report any issue encountered when running the respective model (some of which might be false positives), and “AICc weight” indicates the cumulative model weight, that is it the proportion of the total amount of predictive power provided by all the models contained in each model. Here, the first model shows the lowest AICc value and carries the highest proportion of the cumulative model weight.

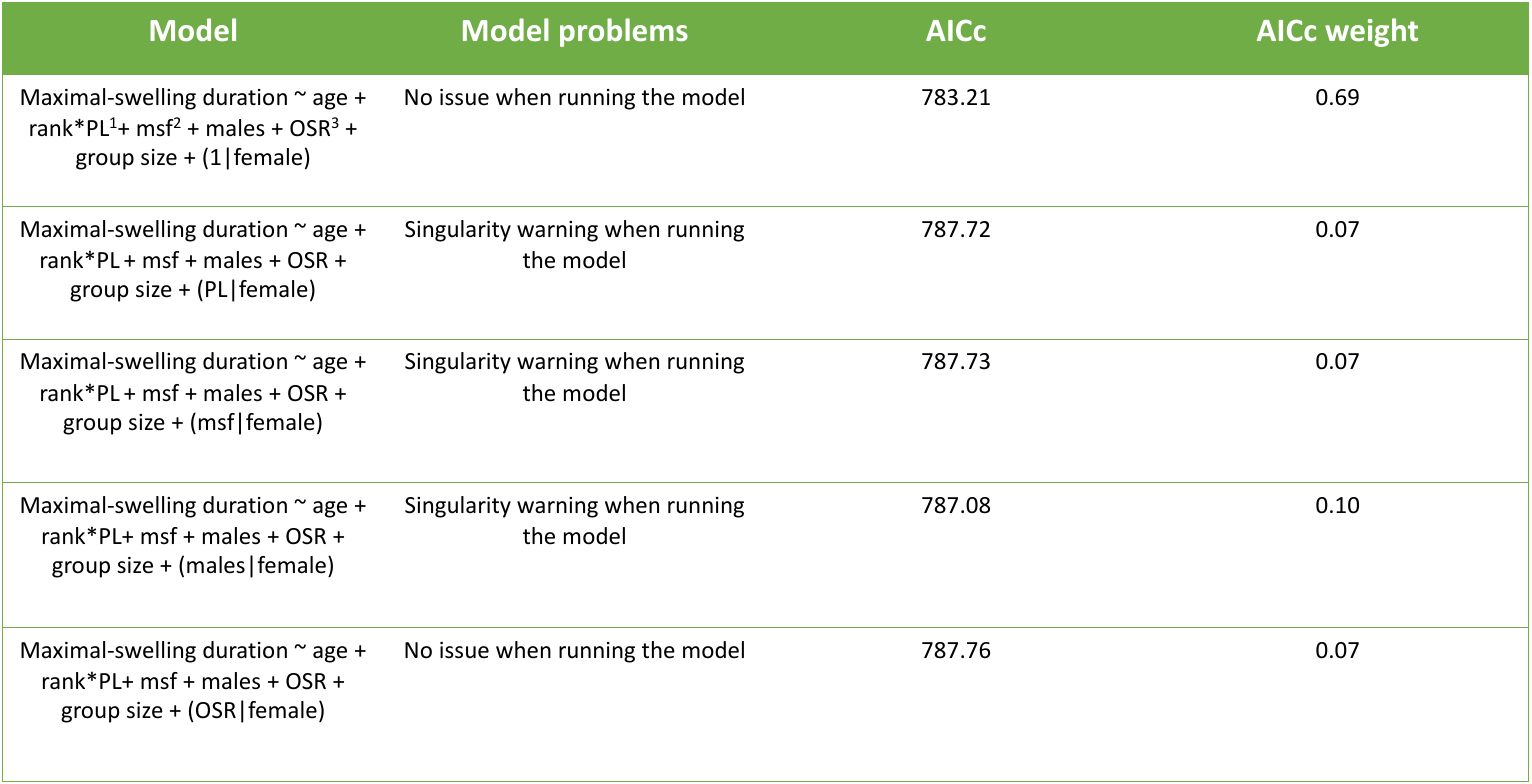

^1^Number of pregnant and lactating females with an interaction with rank; ^2^Number of maximally-swollen females when the focal female started swelling; ^3^Operational sex-ratio as the number of the number of sexually active females to adult males.

**Table A4**

Comparison of our full model to the null model with a likelihood ratio test. The results indicate significant evidence for the two compared models deferring in their explanatory power, as the p-value is greater than our threshold level (0.05). Also, the model with the lower negative log-likelihood value, which in both cases is our selected model, is a better fit.

| Full model | Null model | Results |
| --- | --- | --- |
| Swelling duration ~ age + rank*P females^1^+ rank*L females^2^ + nnmsf^3^ + males + OSR^4^ + (1\|troop/female) | Swelling duration ~ age + rank + (1\|troop/female) | LogLik full model = -453.19  LogLik null model = -460.37  p-value = 0.04516 |
| Maximal-swelling duration ~ age + rank*PL^5^+ msf^6^ + males + OSR + group size + (1\|female) | Maximal-swelling duration ~ age + rank + (1\|female) | LogLik full model = -379.58  LogLik null model = -389.58  p-value = 0.002785 |

^1^Number of pregnant females; ^2^Number of lactating females; ^3^Number of not-maximally-swollen females when the focal starts swelling; ^4^Operational sex ratio; ^5^Number of pregnant and lactating females; ^6^Number of maximally-swollen females when the focal female enters its swelling period.

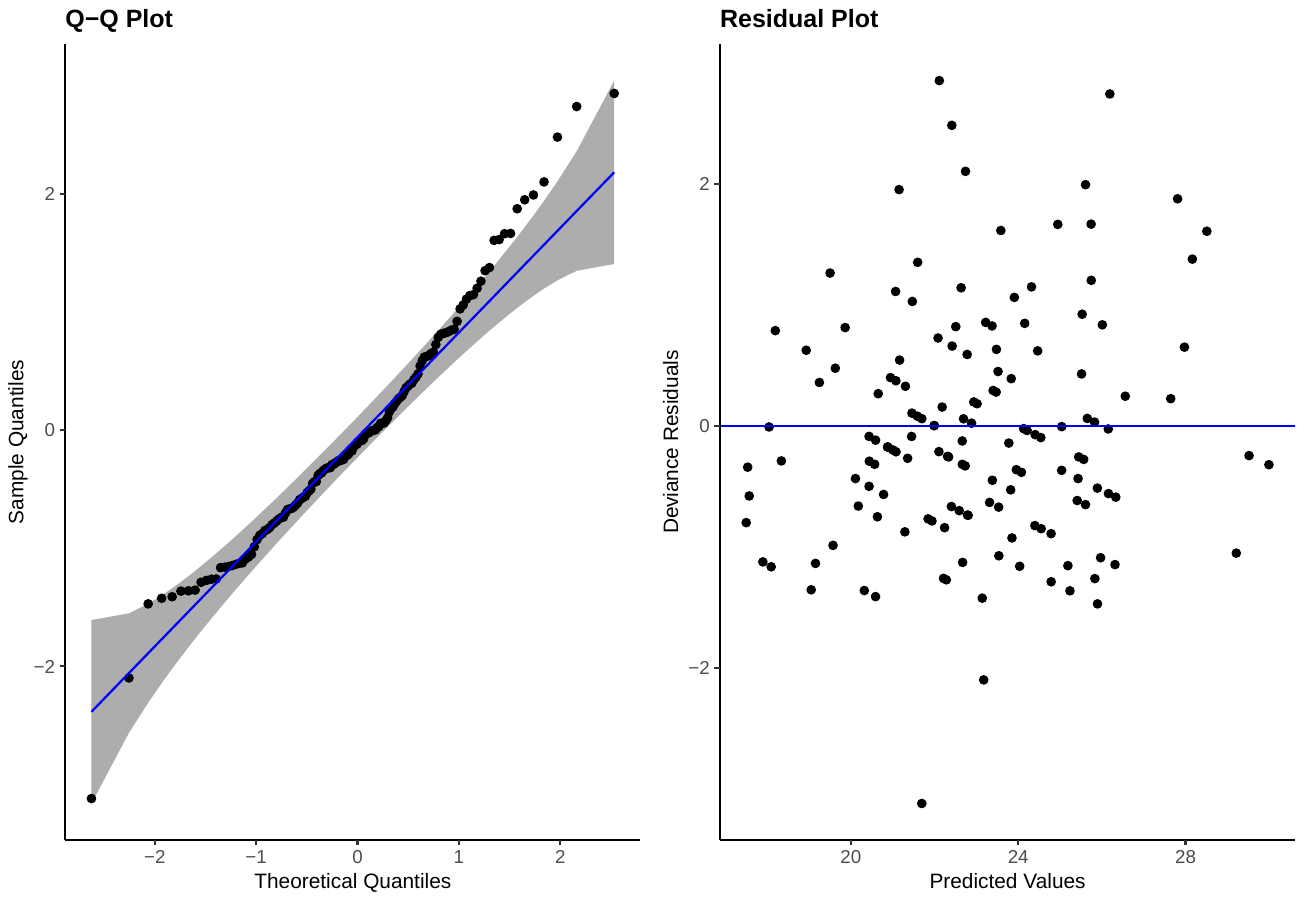

**Figure A1:** Diagnostic plots of swelling duration model to explore if the model fits well the data and if residuals (deviance residuals are plotted) are randomly distributed.

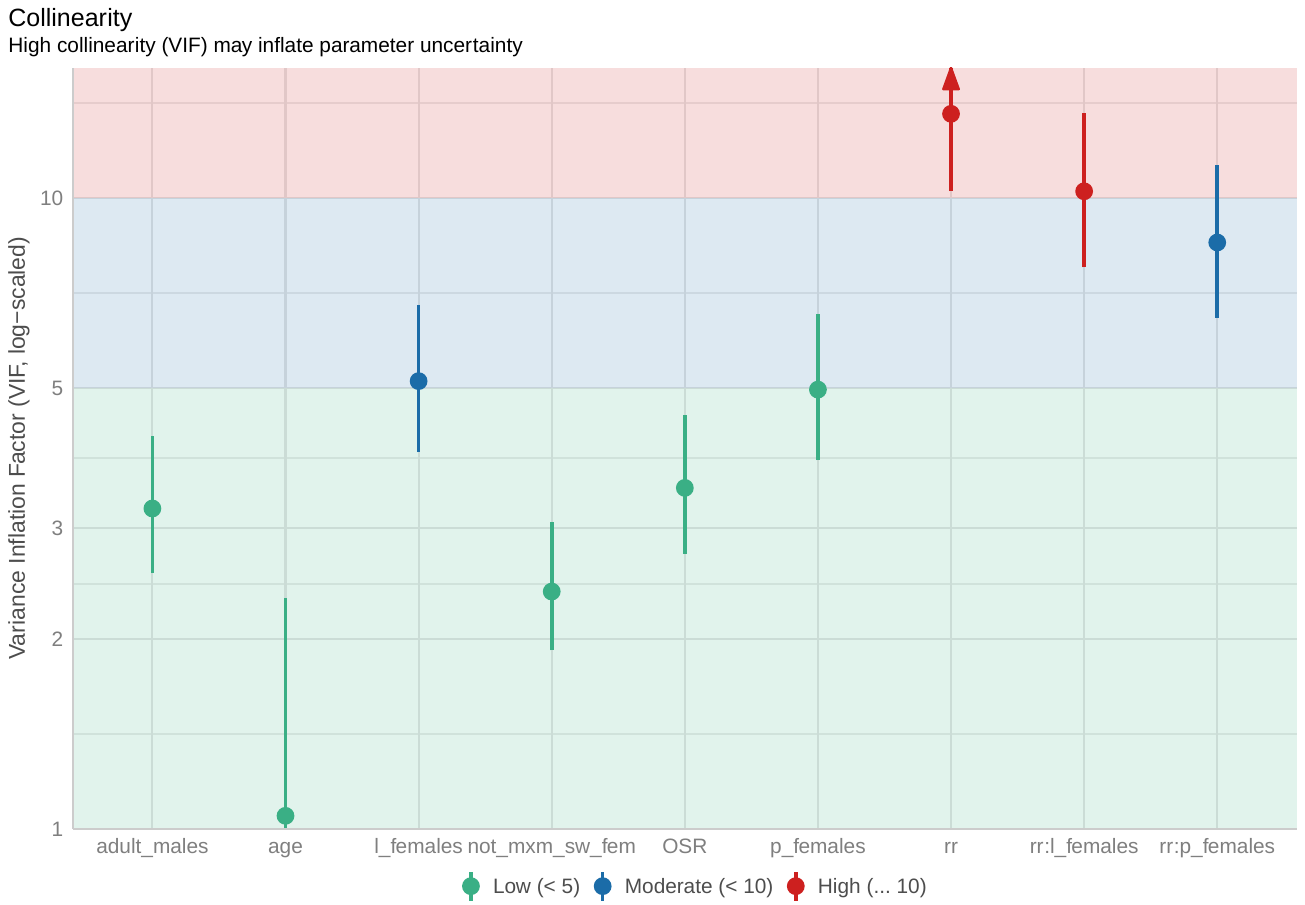

**Figure A2:** Variables of model 1 (swelling duration) show a low (< 5) magnitude of multicollinearity.

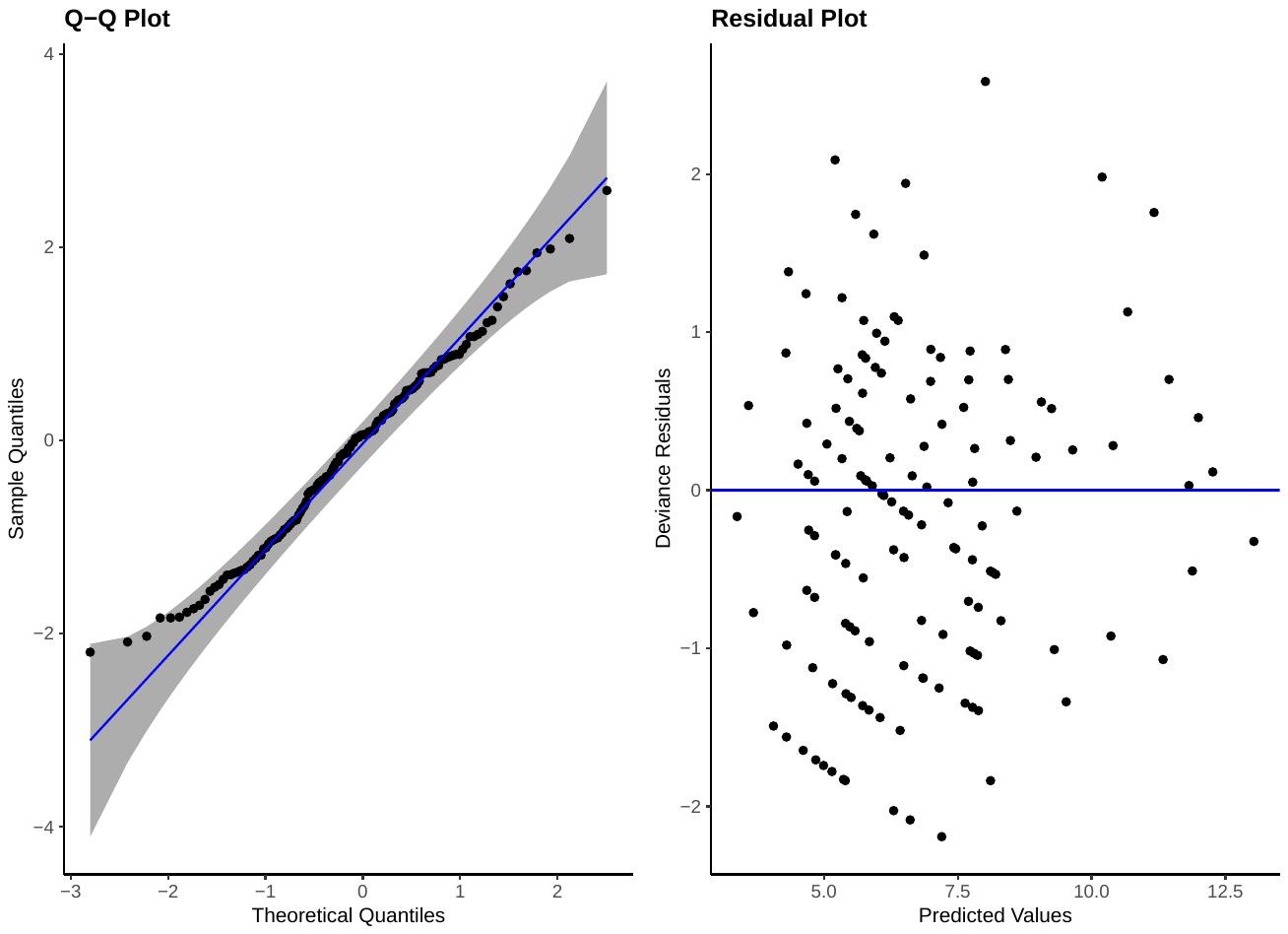

**Figure A3:** Diagnostic plots of maximal-swelling duration model to explore if the model fits well the data and if residuals (deviance residuals are plotted) are randomly distributed.

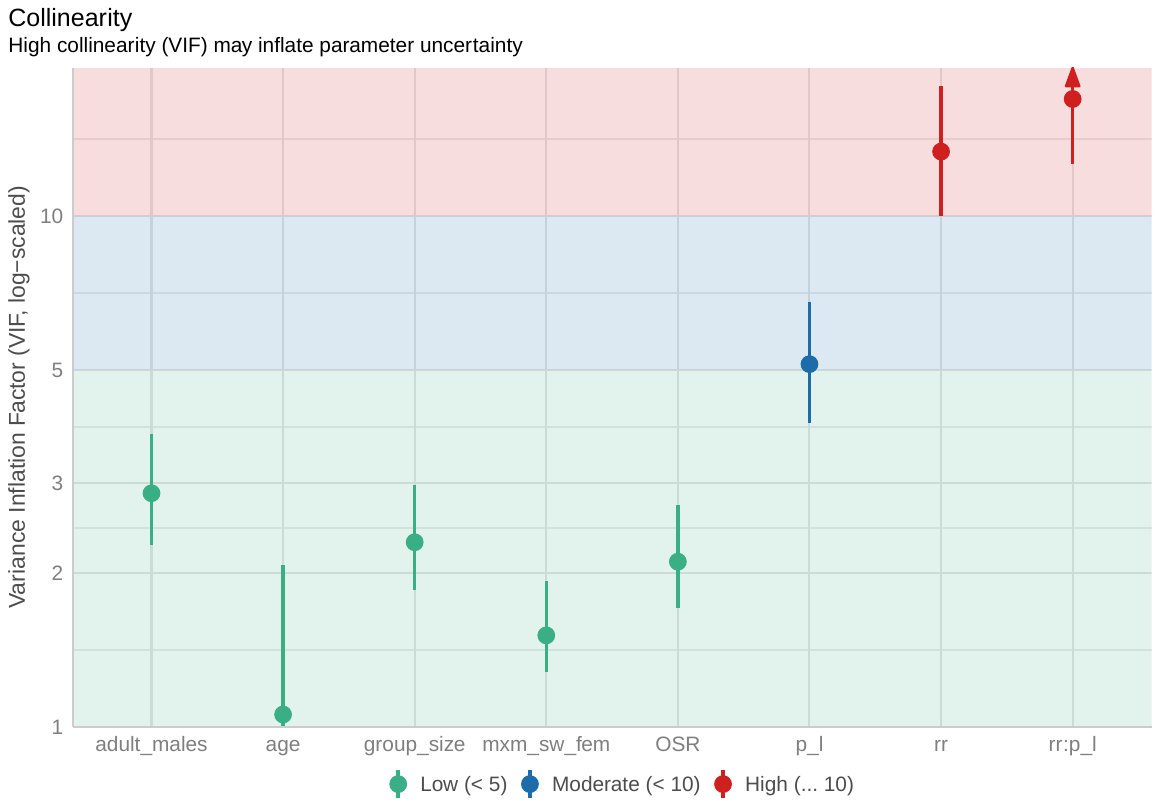

**Figure A4:** Variables of model 2 (maximal-swelling duration) show a low (< 5) magnitude of multicollinearity.

**Table A5**

Repeatability estimates at the level of the female with swelling and maximal-swelling as the dependent variables. Shown are the estimates (R), the proportion of variance of the response variable explained by fixed effects (F), the standard error of the repeatability estimate (SE), and the confidence interval (CI).

| *Response Variable* | *Fixed effects* | *Random effect(s)* | *R* | *F* | *SE* | *CI* | *p-value* |
| --- | --- | --- | --- | --- | --- | --- | --- |
| Swelling duration | age + rank | female ID + troop | 0.036 | 0.051 | 0.049 | [0, 0.167] | 0.251 |
| Maximal-swelling duration | age + rank | female ID + troop | 0.054 | 0.061 | 0.058 | [0, 0.2] | 0.283 |

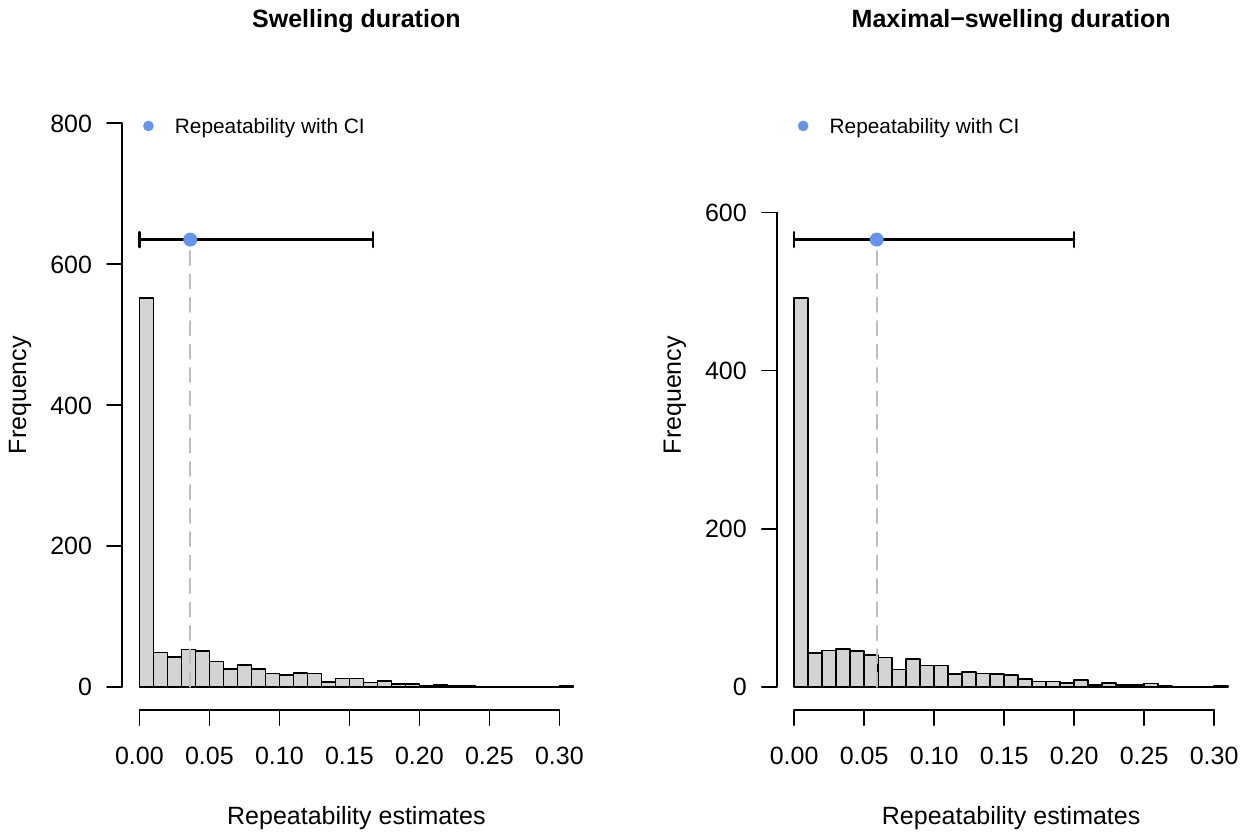

**Figure A5**: Repeatability analysis of the swelling and maximal-swelling duration from 1000 bootstraps (nboot = 1000) at the level of female: rank and age are fixed effects in both models, female ID and troop random effects for swelling and maximal-swelling duration. CI=confidence interval. Original-scale repeatabilities are shown.

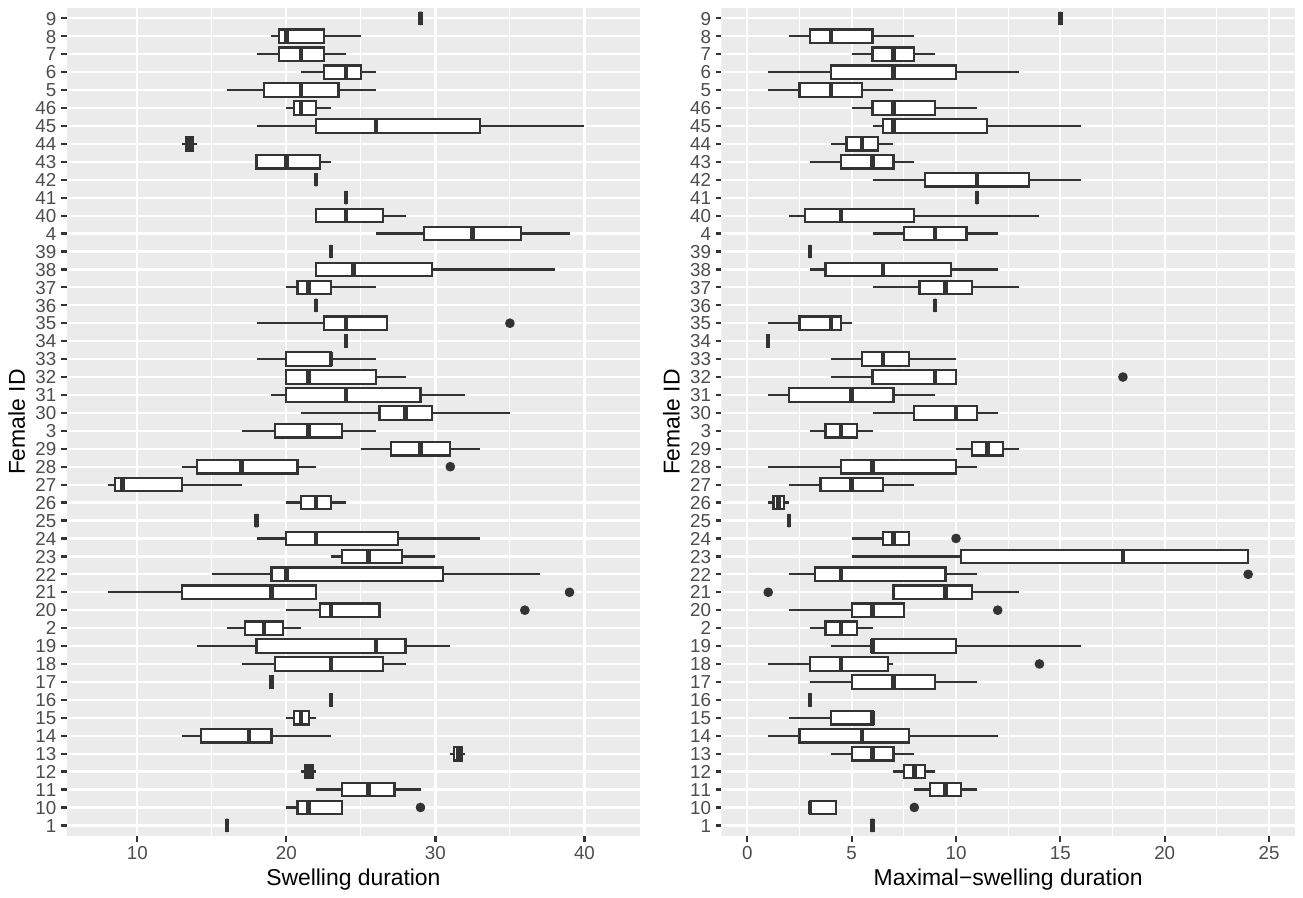

**Figure A6**: Boxplots indicating between and within female variation in swelling (left) and maximal-swelling (right) duration. Outliers are expressed as black dots, with vertical black lines depicting a sole measurement for the respective female. Black lines within the boxes represent the median.

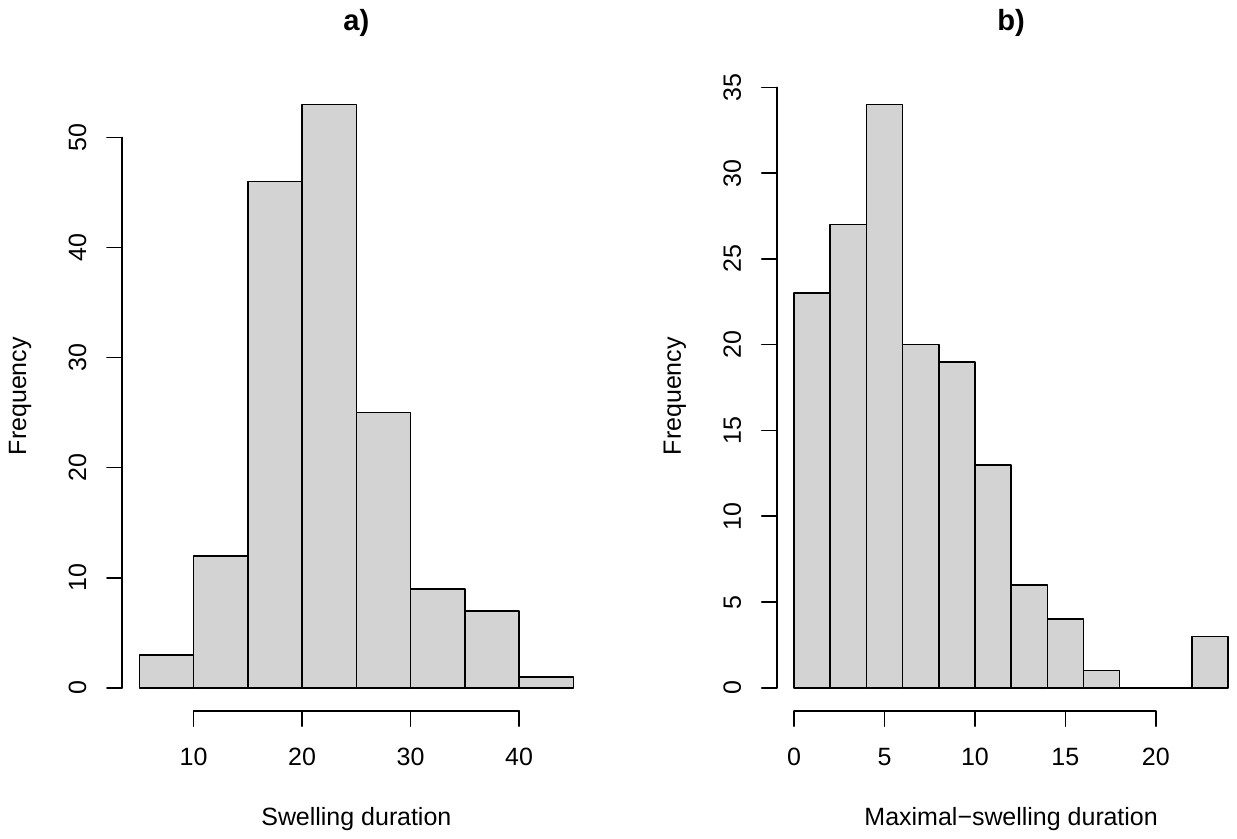

**Figure A7**: The distribution of swelling duration (a) and maximal-swelling duration (b) in this study [number of females a), b) = 48].
